# DaphnAI: A Deep Learning Approach for High-Throughput Zooplankton Community Analysis

**DOI:** 10.1101/2025.07.30.667622

**Authors:** Stylianos Mavrianos, Vera van Santvoort, Sven Teurlincx, Steven AJ Declerck, Kathrin A Otte

## Abstract

Zooplankton are critical components of freshwater ecosystems, mediating energy transfer and regulating trophic dynamics. Monitoring their community composition and population structure is essential for understanding ecological responses to environmental change. However, conventional approaches rely on manual processing and classification, which are time-consuming, prone to error, and unsuitable for high-throughput applications. Here, we present a deep learning framework based on the YOLOv12n-seg architecture for the automated identification, instance segmentation, and quantification of zooplankton from high-resolution images. Trained on mesocosm data containing multiple species zooplankton taxa, our model achieved a mean average precision (mAP@50) of 0.899 and demonstrated a 230-fold speed increase over manual annotation, even on standard non-GPU hardware. Unlike previous approaches that focus solely on classification, our model produces pixel-precise segmentation masks, enabling accurate estimation of individual size metrics. This expands the scope of automated zooplankton monitoring from presence/absence data to demographic assessments, including shifts in size distributions and the possibility to be used for unraveling morphological adaptations. Our work demonstrates how deep learning can overcome longstanding limitations in ecological monitoring by enabling scalable, reproducible, and high-resolution quantification of community composition. This approach opens the door to large-scale, long-term studies of freshwater ecosystems.

## 1. Introduction

Understanding species composition dynamics and population size structure is essential for elucidating ecological processes and predicting community responses to environmental change. Freshwater ecosystems, in particular, are highly sensitive to such changes and are increasingly impacted by stressors like climate change, pollution, and eutrophication (Dudgeon et al., 2006; Heino et al., 2009; Woodward et al., 2010). Within these systems, zooplankton play a pivotal role in transferring energy through food webs and regulating primary production (Ebert, 2005; Lampert, 1987; Sterner, 2009). Among them, *Daphnia* stand out as keystone species and are widely used model organisms in ecology, evolution, and ecotoxicology due to their small size (∼1–5 mm), short generation times, and high sensitivity to environmental stressors (Ebert, 2005).

Given their ecological importance, accurately monitoring zooplankton populations and communities is essential. Traditionally, methods have relied on manual measurement techniques. Manual counts and species classification from images remain common practice in many studies (Adoteye et al., 2015; McKnight et al., 2023). While effective in controlled conditions and small sample sizes, such methods suffer from inherent limitations. Manual counting and classification is time-consuming, prone to observer bias, and difficult to scale, particularly in long-term or high-throughput experimental designs. Subsampling approaches, which extrapolate population-level properties from smaller samples, offer time savings (Milbrink and Bengtsson, 1991) but can introduce significant errors (Erickson et al., 2014). The demand for more objective and efficient data collection has led to the adoption of semi-automated image analysis methods. These include classical computer vision approaches such as thresholding, edge detection, and shape fitting algorithms (Becker et al., 2022; Færøvig et al., 2002; Mallard et al., 2013). However, while such methods can improve throughput to some extent, they typically lack robustness. Slight changes in lighting, focus, or background noise can lead to substantial inaccuracies, particularly when dealing with translucent or overlapping organisms. Moreover, studies have shown that image-based techniques can systematically underrepresent true population sizes when compared to manual counts (Hooper et al., 2006; Liess et al., 2006). They also typically require specialized and expensive hardware setups (Duckworth et al., 2019).

In recent years, deep learning has significantly advanced image-based analysis in many fields, including ecology. Convolutional neural networks (CNNs) have proven highly effective in image classification. These models learn features directly from data, eliminating the need for manual feature design (O’Shea and Nash, 2015). They can efficiently analyze large datasets, reducing manual effort and improving reproducibility. For example, Karatzas et al., 2020 used CNNs to detect malformations in *Daphnia magna* exposed to nanomaterials, automating complex toxicological assessments. Kim et al., 2023 developed a machine learning-based video tracking system to monitor *Daphnia* behavior for rapid ecotoxicity testing. Similarly, Ma et al., 2024 applied deep learning techniques to accurately count and classify adult and neonate *Daphnia*, achieving results comparable to manual annotation. *LeDaphNet* (Rutter et al., 2024) used a CNN in combination with a standard scanner and 3D-printed vessel to semi-automate population counts of *D. magna*, achieving over 92% accuracy. Recent reviews further highlight that deep learning has enabled not only detection and classification but also behavioral analysis and ecological modeling for both phyto– and zooplankton, accelerating data acquisition and reducing human bias in plankton ecology (Bachimanchi et al., 2024). However, despite their success in detection and classification tasks, these approaches are generally restricted to identifying and, in some cases, localizing objects within an image, but they lack the capability for instance segmentation. They cannot accurately delineate and separate each individual organism by outlining its precise boundaries within complex or crowded scenes.

Among the most promising deep learning architectures for object detection and segmentation is the YOLO (You Only Look Once) family of models. YOLO models employ a single-stage object detection framework, making them both computationally efficient and accurate. Since the original YOLOv1 model was introduced in 2016 (Redmon et al., 2016), successive versions have integrated state-of-the-art components such as spatial pyramid pooling, cross-stage partial networks, attention modules, and multi-scale feature extraction (Sapkota et al., 2024). The most recent version, YOLOv12 (Tian et al., 2025), incorporates advanced transformer-based modules (Vaswani et al., 2017) and supports both object detection and instance segmentation. This functionality allows for not only the localization of objects via bounding boxes but also the generation of pixel-precise segmentation masks, which in turn allows identification, counting, as well as the measurement of surface area and other metrics that could serve as proxies for body size, crucial for assessing shifts in size distributions.

Recent studies have demonstrated the potential of YOLO-based models for monitoring small aquatic organisms in complex environments to answer ecological questions. YOLO models enable real-time tracking of aquatic organisms, as demonstrated by their application in monitoring the swimming behavior of *D. magna* exposed to mercury chloride (Qin et al., 2024). Their strength in detecting objects in cluttered backgrounds and under varying lighting conditions has been critical for tasks such as identifying invasive species like water hyacinth in natural aquatic environments and classifying benthic species from deep-sea crawler footage, reducing the need for manual sorting and improving the scalability of ecological surveys (Ji et al., 2025; Ortenzi et al., 2024). Furthermore, YOLO’s ability to detect small objects with high accuracy makes it especially suitable for fine-scale ecological monitoring, such as locating *Pomacea canaliculata* eggs in noisy agricultural scenes (Huang et al., 2023)

In this study, we introduce the first application of a YOLO-based model for automated identification, counting, and instance segmentation of *Daphnia* from high-resolution images. Our model is designed to operate on images containing multiple species of *Daphnia*. Unlike existing methods, it does not rely on manual pre-filtering or idealized imaging conditions. Instead, it is trained to recognize and segment individuals under variable lighting conditions, with overlapping individuals and complex organic backgrounds. By applying this model to large datasets from mesocosm experiments, we demonstrate its capacity for high-throughput analysis of zooplankton community composition and size distribution. This approach offers a scalable tool for ecological monitoring, supporting research into the dynamics of zooplankton populations and their role in ecosystem processes.

## 2. Materials and Methods

### 2.1 Model training

#### 2.1.1 Image Acquisition and Preprocessing

The analysis was performed on samples of a mesocosm experiment consisting of 48 outdoor water tanks (volume: 180L). The experiment consisted of three experimental groups, with 16 single-species mesocosms per group. The treatments consisted of either *Daphnia galeata*, *Daphnia pulex*, or *Simocephalus vetulus*. From each group, zooplankton samples were taken from 6 randomly selected mesocosms with a tube sampler, concentrated with a net (mesh size: 80 µm) and preserved with Lugol’s Iodine solution (5%). Each sample was scanned twice, using an Epson Perfection V850 Pro Flatbed scanner. The first time samples were scanned in their fixative, the second time after being washed with tap water over a screen of 80 µm. This was done to introduce variability in background conditions, helping the model learn to focus on meaningful features and improve its ability to detect target objects across variable visual settings.

The final dataset consisted of 36 high-resolution images (see Figure 1). Due to their large size, the images were systematically cropped to improve computational efficiency. Each image was divided into 77 chunks of 1024 × 1024 pixels. This pre-processing resulted in 2.772 chunk images, which were used for training, validation, and testing.

**Figure 1:**
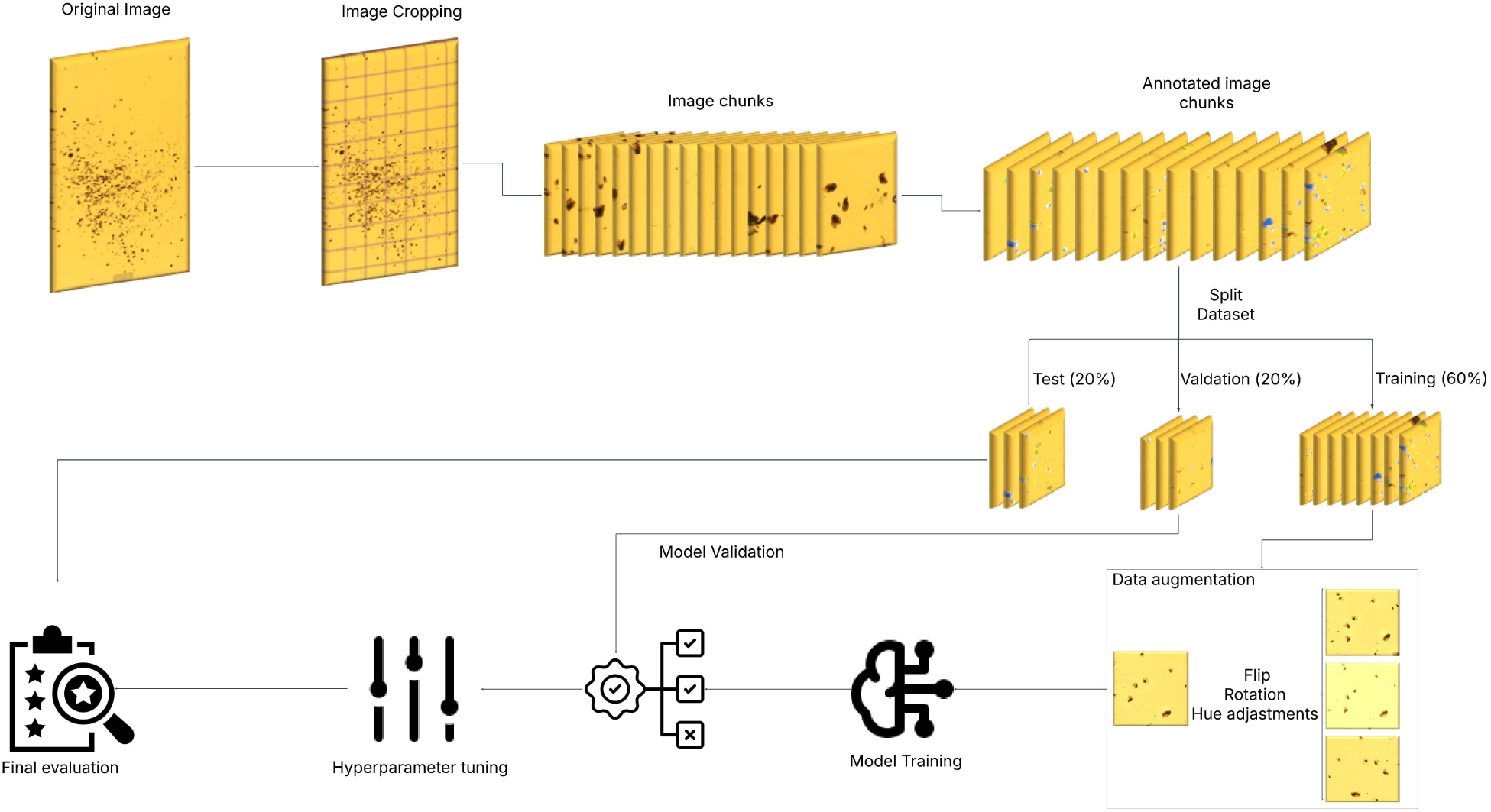
Image analysis pipeline for training the object detection and segmentation model. The process begins with the original image, which is cropped into smaller overlapping image chunks. These chunks are manually annotated and then split into training (60%), validation (20%), and test (20%) sets. The training set undergoes data augmentation (e.g., flipping, rotation, brightness adjustments) to enhance model robustness. Model training is followed by validation and hyperparameter tuning. Once optimized, the model undergoes final evaluation using the independent test set.

#### 2.1.2 Image Annotation

Ground-truth annotations (see Figure 2), which serve as the definitive reference for training and evaluating machine learning models, were generated using a combination of manual and semi-automatic labeling. The semi-automatic annotations were facilitated by the Segment Anything Model 2 (SAM2; Ravi et al., 2024) which enabled precise segmentation of objects and significantly reduced manual workload. However, in cases where SAM2 failed to produce accurate results, manual annotation was performed to ensure high-quality labels. In this context, a ground-truth annotation consists of two key components: (i) a segmentation polygon mask, which outlines the exact pixel-wise boundaries of an object in the image, and (ii) a class label, which assigns the object to one of the predefined categories (see SFigure 1a). These detailed annotations provide the model with explicit information about what the object is and where it is located within the image, enabling supervised learning for object detection and segmentation tasks. Seven morphologically distinct classes were identified and labeled: *Daphnia pulex, Daphnia galeata, Simocephalus vetulus*, ballooned *Daphnia*, Cyclopoid copepod, parthenogenetic eggs (either free or retained inside *Daphnia*), and ephippia (either free or contained within *Daphnia*). The term ballooned *Daphnia* refers to individuals exhibiting an atypically swollen body morphology, caused by the preservation method. To guide consistency, we classified individuals as ballooned when they no longer exhibited the typical Daphnia silhouette, such as a well-defined head, tail spine, and body curvature (see SFigure 2). In addition to the three primary experimental classes, we included copepods because of their frequent occurrence in the samples, and ephippia along with parthenogenetic eggs as important indicators of reproductive activity and fitness.

**Figure 2:**
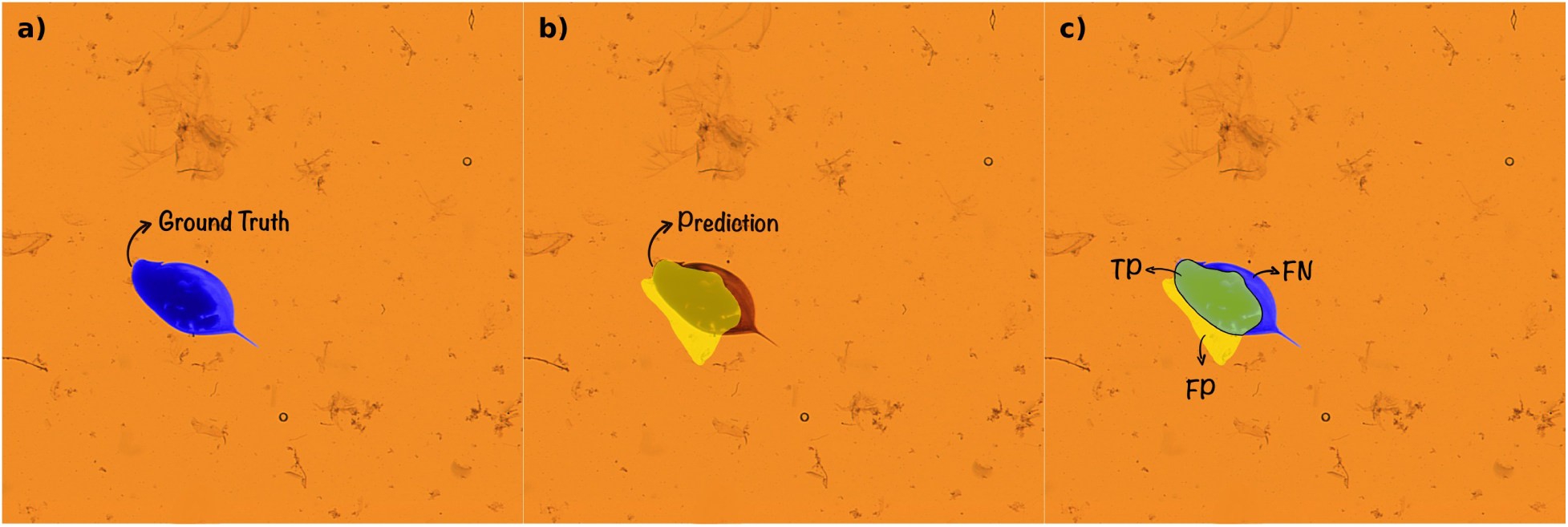
Visualization of instance segmentation prediction accuracy using Intersection over Union (IoU) metrics. a) Ground truth mask (blue) showing the annotated Daphnia individual. b) Model prediction mask (yellow) overlaid on the same image. c) Segmentation performance breakdown: True Positive (TP, green) indicates the overlapping region between ground truth and prediction; False Negative (FN, blue) shows the part of the ground truth missed by the model; False Positive (FP, yellow) represents the area falsely predicted outside the true object boundary.

#### 2.1.3 Dataset Partitioning and Augmentation

Following annotation, the dataset (the 2.772 annotated images) was stratified into three subsets with 60% allocated for training, 20% for validation, and 20% for testing. The training set was used to teach the model how to detect, identify, and segment features, while the validation set helped fine-tune the model during training, performance validation, and hyperparameter tuning by evaluating how well it generalizes to data it has not directly learned from. The test set, which the model had never seen before, was reserved for final performance evaluation. To enhance model robustness and mitigate overfitting, data augmentation techniques (Shorten and Khoshgoftaar, 2019; Yang et al., 2023) were applied only to the training set using the Albumentations python library (Buslaev et al., 2020). Overfitting occurs when a model memorizes the training data, including noise and irrelevant patterns, rather than learning the underlying general features that enable it to perform well on new, unseen data. To counter this, augmentations introduced variability into the training data, helping the model learn more generalizable patterns. These augmentations included geometric transformations such as horizontal and vertical flipping, random rotations, and photometric adjustments involving hue, brightness, and saturation adjustments. Additionally, Mixup and Mosaic augmentation were used to increase dataset diversity. Mixup augmentation generates synthetic training samples by blending two images and interpolating their corresponding labels, thereby enhancing generalization. Mosaic augmentation, on the other hand, constructs a composite image by merging four distinct images, providing the model with complex contextual variations that improve its ability to detect and segment objects in diverse conditions.

#### 2.1.4 Model Training and Performance Monitoring

A custom YOLO12n-seg (version 12 nano segmentation) model, a computationally efficient, attention-centric deep learning architecture tailored for instance segmentation, was trained specifically for this task. The model was initialized without pretrained weights, allowing for full customization to the unique characteristics of our dataset. Training was conducted for up to 1000 epochs (i.e., complete passes through the entire training dataset) with early stopping applied, using a patience parameter of 100 epochs. This means that training was terminated once no significant improvement was observed on the validation set for 100 consecutive epochs. The best model performance was achieved at epoch 508, after which continued training did not yield further improvements. The training was performed on a single NVIDIA H100 GPU with a batch size dynamically based on 80% of the maximum available GPU memory. The Adam optimizer was used with a base learning rate of 0.01. The entire training process lasted 4 hours and 58 minutes.

To evaluate the learning progress of the model during training, we monitored the evolution of two key loss functions: classification loss and segmentation loss. These loss functions quantify the discrepancy between the model’s predictions and the ground-truth data, serving as objective measures of model performance. The classification loss assesses how accurately the model assigns class labels to detected individuals. A lower classification loss indicates that the model is increasingly confident and correct in identifying the appropriate class for each segmented object. The segmentation loss evaluates the model’s ability to delineate the spatial boundaries of each individual. It measures how closely the predicted segmentation polygon masks align with the manually annotated ground truth polygon masks. High segmentation loss suggests that the predicted outlines deviate significantly from the true shape, whereas a declining segmentation loss over time reflects more precise and accurate spatial predictions.

In order for a model to actively learn features, both loss functions should progressively decrease as the training proceeds, not only on the training data but also on the validation. The training loss function reflects how well the model performs on the data it sees during training. In contrast, the validation loss function indicates how well the model performs on unseen data and it works as a proxy for real-world performance. If the training loss continues to drop but the validation loss stalls or increases, it may indicate overfitting, where the model is memorizing the training data rather than learning generalizable patterns.

### 2.2 Model Performance Metrics

Model performance evaluation was done using standard evaluation metrics: precision (see Equation 1), recall (see Equation 2), F1-score (see Equation 3), and mean Average Precision (mAP; see Equation 4) at both mAP@50 and mAP@50–95. Precision quantifies the proportion of correctly predicted objects (true positives) out of all instances predicted as belonging to that class. Recall measures the proportion of correctly predicted objects out of all ground-truth instances for that class. The F1-score provides a balanced measure of the model’s ability to minimize both types of error. The mAP is a standard evaluation multi-class metric in instance segmentation that reflects both how accurately a model classifies objects (e.g., *Daphnia pulex*) and how precisely it localizes them within an image. It is based on the calculation of Average Precision (AP; see Equation 5). For each object class, two main components need to be considered. The first is the confidence score, which reflects how certain the model is that a particular object is present in a specific part of the image. The second component is the Intersection over Union (IoU; see Equation 6). This is a way of measuring how well the model’s predicted outline of the organism overlaps with the actual, correct annotation (ground truth; see Figure 2). An IoU of 1 means a perfect match; an IoU of 0 means no overlap at all (see SFigure 1b). Thus, a prediction is counted as a true positive only when the overlap (IoU) exceeds a certain predefined threshold, such as 0.50 (meaning at least 50% overlap), and when the confidence score is above a minimum predefined level. If either of these conditions is not met, the prediction might be counted as a false positive or ignored (false negative). Thus, the AP is computed as the area under the precision–recall curve (AUC), generated by varying the confidence threshold from 1 to 0. The mAP is then obtained by averaging AP values across all classes. The mAP@50 refers to the mean Average Precision at an IoU threshold of 0.50, while mAP@50–95 provides a stricter evaluation by averaging AP scores across IoU thresholds from 0.50 to 0.95 in steps of 0.05 for all classes.

#### 2.2.1 Model Validation

Model validation was performed on the validation set using a fixed Intersection over Union (IoU) threshold of 0.70, corresponding to AP@70. The PR curves, which visualize the trade-off between false positives and false negatives across different decision thresholds, help assessing the model’s ability to maintain a balance between precision and recall. As part of post-training hyperparameter tuning, F1– confidence plot was used to identify the confidence threshold that best balances precision and recall for each object class.

#### 2.2.2 Model Evaluation

Final model evaluation was conducted on the test set, which was kept entirely separate from training and validation to ensure an unbiased assessment of generalization performance. Using the tuned confidence threshold determined during validation, we computed key metrics including mAP at IoU thresholds of 0.50 and 0.50–0.95, as well as class-specific precision, recall, and F1-score. A normalized confusion matrix was also generated to assess prediction accuracy and misclassification patterns.

## 3. Results

### 3.1 Model Training

The segmentation loss exhibited a very strong downward trend followed by a further gradual decline throughout training, indicating effective learning by the model (see Figure 3a). The validation segmentation loss closely followed the training curve, suggesting no overfitting. Similarly, classification loss decreased over time, with the validation classification loss tracking the training loss closely (see Figure 3b). These patterns indicate strong generalization and stable learning. Early stopping was triggered at epoch 508, the point at which no further improvement in validation loss was observed, ensuring the model achieved an optimal balance between learning and overfitting prevention.

**Figure 3:**
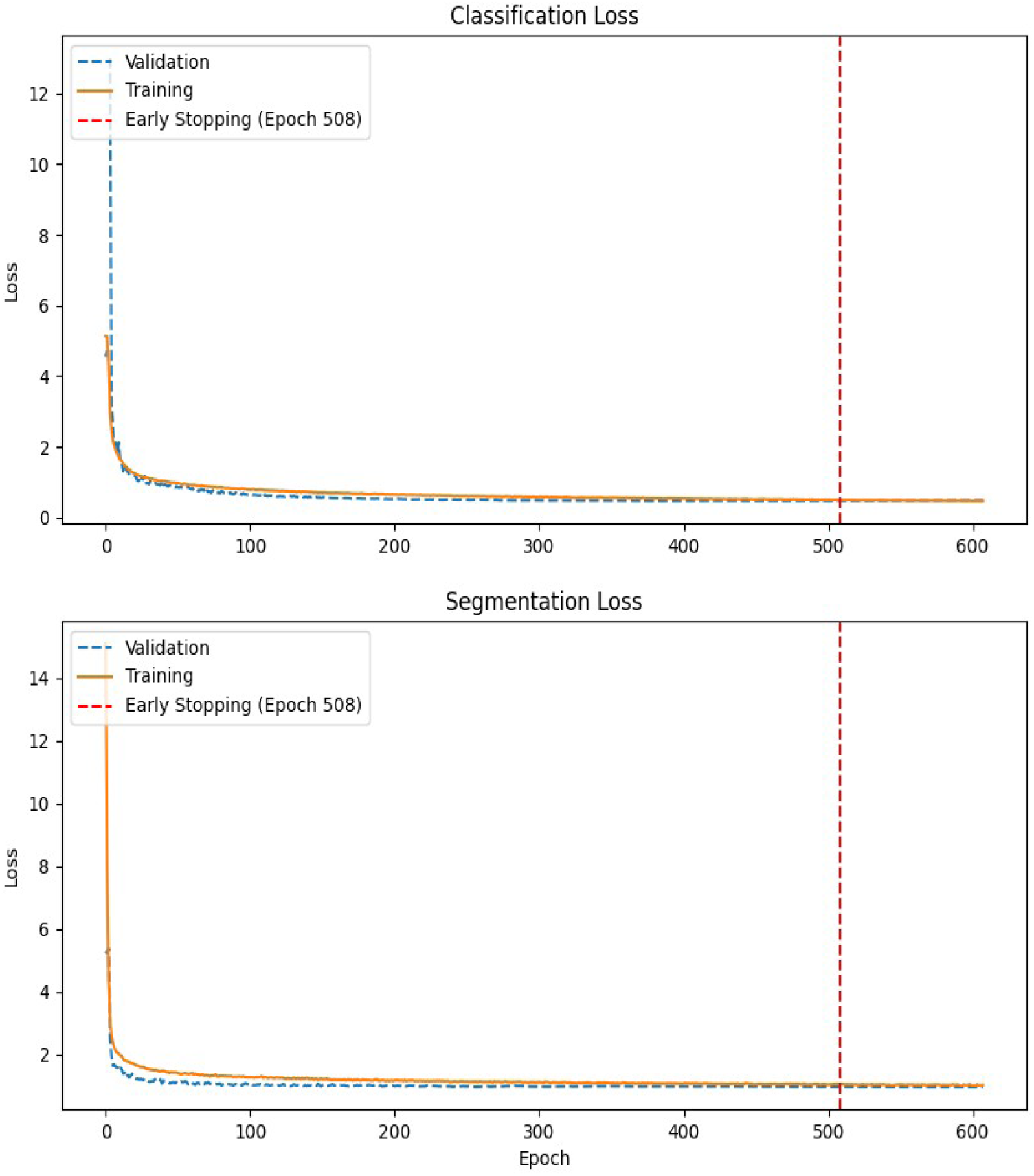
Illustration of the loss curves over the training epochs, comparing training and validation losses for segmentation and classification, respectively. a) shows the segmentation loss over training epochs, with the x-axis representing the number of epochs and the y-axis the segmentation loss value. b) shows the classification loss across training, with the same axes representing epochs and classification loss, respectively. In both plots, training and validation losses are shown to assess convergence and generalization. A dashed vertical line at epoch 508 marks the early stopping point, indicating the moment when training was automatically halted due to a plateau in validation performance, thereby preventing overfitting.

### 3.2 Model Validation

The PR curves revealed high average precision for most classes, with *D. galeata* (0.968), *S. vetulus* (0.964), and *Ballooned* (0.957) showing particularly strong performance (Figure 4a). The F1–Confidence curve showed that the model reached peak performance (F1 = 0.88) at a confidence threshold of 0.377. This threshold was selected as part of a post-training hyperparameter tuning step aimed at maximizing classification performance. Most classes maintained high F1-scores across a wide range of thresholds (see Figure 4b).

**Figure 4:**
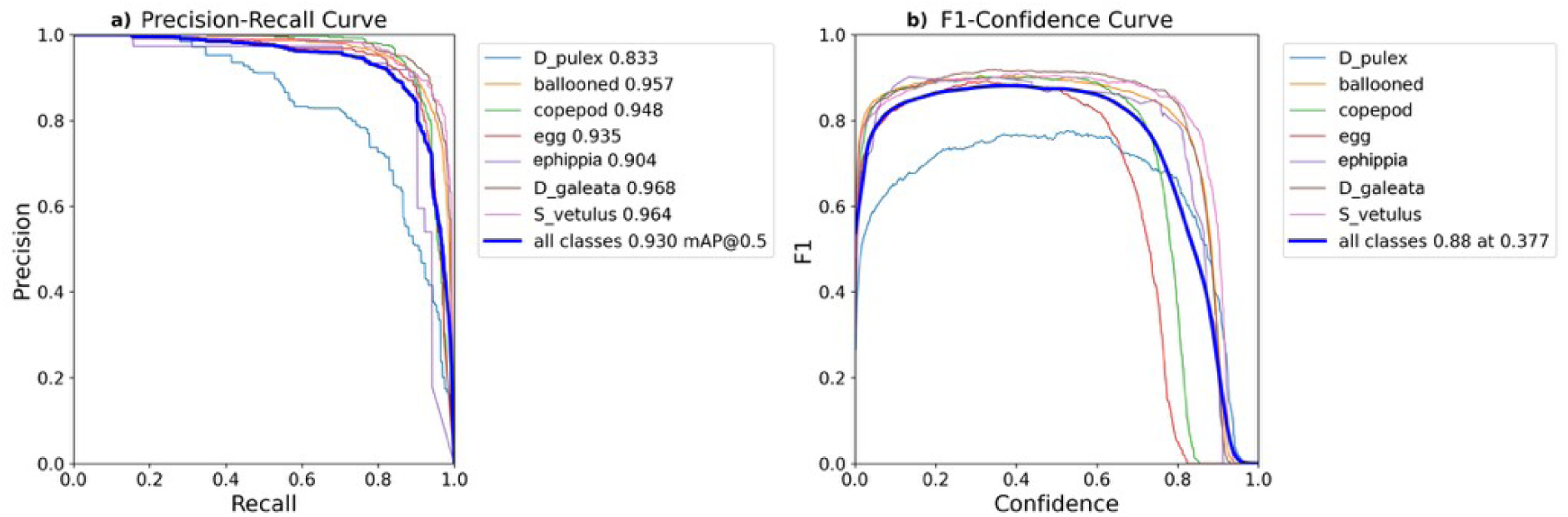
a) Precision–Recall (PR) curves for each annotated class. These curves illustrate the trade-off between precision and recall at varying confidence thresholds (1 to 0). Each line represents a specific class, with curves closer to the upper-right corner indicating stronger detection performance. The area under each curve corresponds to the class-specific average precision (AP). The thick blue line shows the macro-averaged performance across all classes. b) F1-Confidence curves for each annotated class. These curves depict the relationship between model confidence thresholds and the resulting F1-score, highlighting the optimal point at which precision and recall are most balanced. The bold blue line represents the macro-averaged F1-score across all classes.

### 3.3 Model Evaluation

The model achieved high classification performance across most categories. *S. vetulus* and ephippia were correctly classified in 95% and 92% of instances respectively, while *D. galeata* reached a correct classification rate of 91%. However, some misclassifications occurred: 14% of *D. pulex*, 5% of *D. galeata*, and 4% of *S. vetulus* instances were incorrectly assigned to the ballooned class. Additionally, 33% of all background misclassifications were falsely labeled as ballooned (see Figure 5). A further 3% of *D. pulex* individuals were misclassified as *D. galeata*; these individuals had significantly smaller surface areas compared to correctly classified *D. pulex* (see Figure 6; p = 0.042, Welch’s t-test).

**Figure 5:**
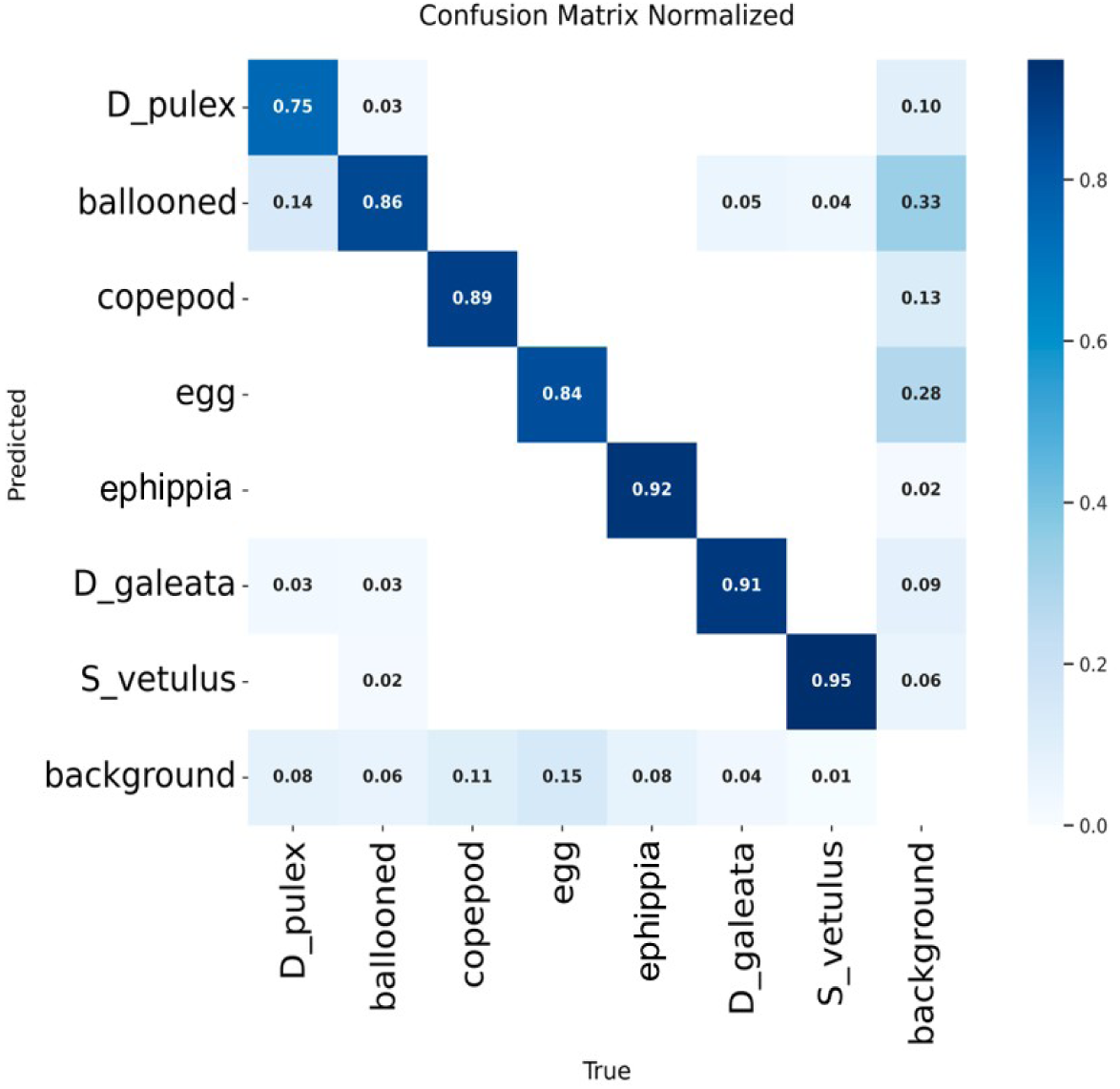
The normalized confusion matrix provides an assessment of the model’s classification performance on the test set of the dataset. The x-axis represents the true class labels (i.e., the ground truth), while the y-axis corresponds to the predicted class labels (i.e., the model’s predictions). Each value in the matrix indicates the proportion of instances of a given true class that were assigned to a particular predicted class, with all values normalized between 0 and 1. High values along the diagonal signify strong classification accuracy (true positives), whereas off-diagonal values represent misclassifications. The background class accounts for regions in the image where no objects of interest are present, yet the model incorrectly assigns them to a category. These cases represent false positives, where the model mistakenly detects an object in areas that should remain unclassified. In the confusion matrix, misclassifications into the background category suggest that the model either failed to recognize an object correctly or falsely classified non-object regions as belonging to a specific class (confidence score was set at 0.377 and IoU at 0.7).

**Figure 6:**
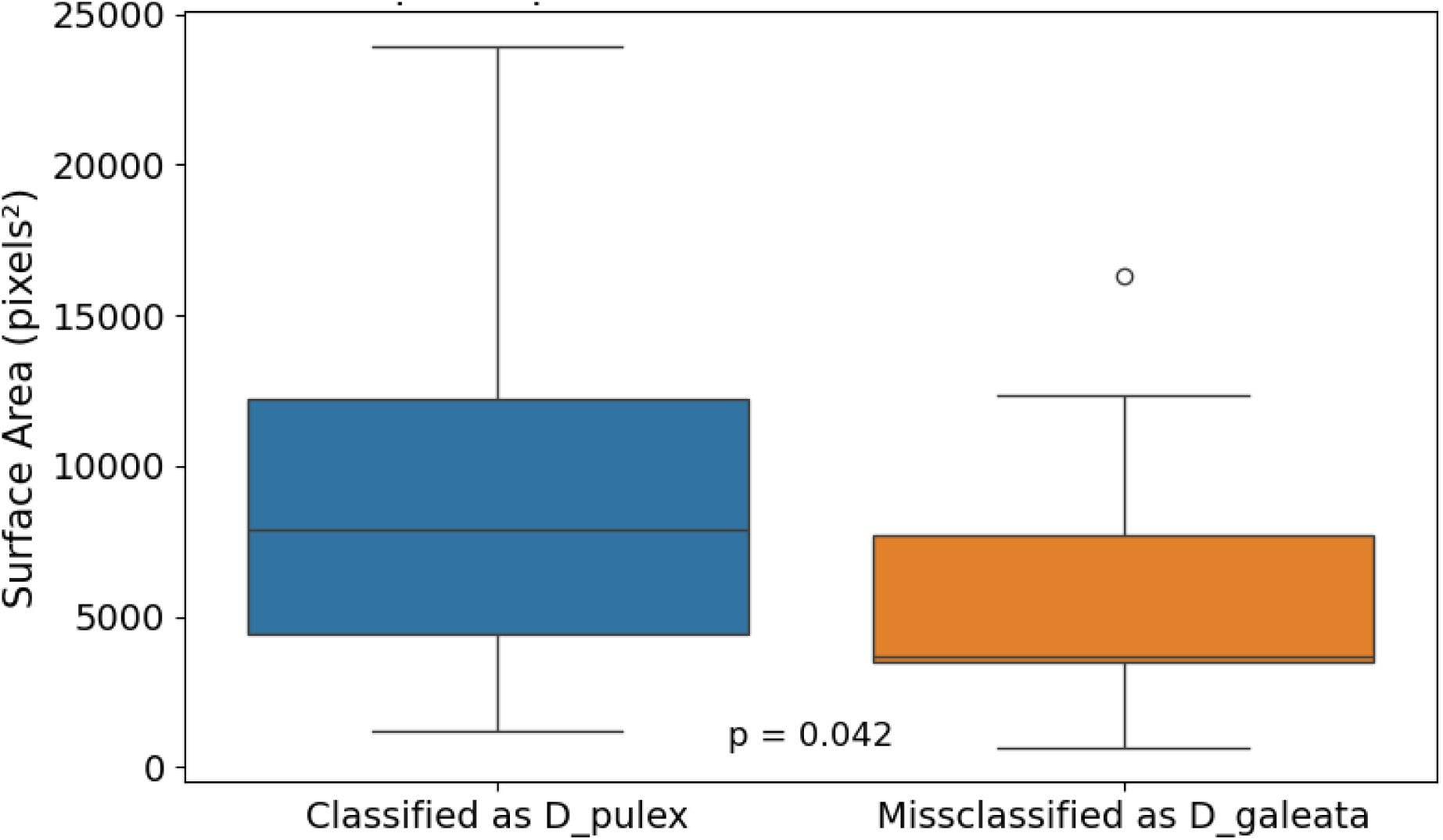
Surface area comparison between correctly classified D. pulex and those misclassified as D. galeata. Individuals misclassified as D. galeata had significantly smaller surface areas (p = 0.042, Welch’s t-test), suggesting a size-dependent classification bias. Surface area is measured in pixels².

Overall, the model achieved a mAP@50 of 0.899 and mAP@50–95 of 0.659, indicating strong generalization across a range of localization accuracies. The best-performing classes included *Cyclopoid copepod* (AP = 0.938), *S. vetulus* (0.928), ephippia (0.927), and *D. galeata* (0.927), all demonstrating high precision, recall, and F1-scores (see Table 1; confidence score was set at 0.377 and IoU at 0.7).

**Table 1:** Segmentation performance of the trained model on the test set. The table summarizes instance segmentation metrics across all annotated classes using mask-based predictions. Reported values include F1-score, precision, recall (confidence score was set at 0.377 and IoU at 0.7), mean Average Precision at IoU threshold 0.5 (mAP@50), and mean Average Precision averaged over thresholds from 0.5 to 0.95 in increments of 0.05 (mAP@50–95). The “all” row represents macro-averaged performance across all classes.

| **Class** | **Precision** | **Recall** | **F1-score** | **mAP50** | **mAP50-95** |
| --- | --- | --- | --- | --- | --- |
| **all** | 0.861 | 0.881 | 0.871 | 0.899 | 0.659 |
| *D_pulex* | 0.718 | 0.781 | 0.749 | 0.777 | 0.671 |
| *ballooned* | 0.922 | 0.869 | 0.895 | 0.92 | 0.783 |
| *copepod* | 0.94 | 0.895 | 0.917 | 0.938 | 0.544 |
| *egg* | 0.851 | 0.829 | 0.84 | 0.878 | 0.379 |
| *ephippia* | 0.889 | 0.923 | 0.906 | 0.927 | 0.724 |
| *D_galeata* | 0.877 | 0.92 | 0.898 | 0.927 | 0.759 |
| *S_vetulus* | 0.826 | 0.949 | 0.884 | 0.928 | 0.752 |

### 3.4 Annotation Efficiency and Inference Speed

A key benefit of the trained YOLOv12n-seg model lies in its ability to dramatically accelerate the annotation process. Manual annotation of the 2.772 chunked training images required approximately 90 hours of manual effort. In contrast, once trained, the model can annotate and extract both species classifications and key anatomical traits from a single full-resolution image, consisting of 77 chunked segments (each 1024 × 1024 pixels), in just 39 seconds. This inference was performed on a standard Dell laptop (Intel Core i5, 12 CPUs, 15 GB RAM) without GPU acceleration. This represents a 230-fold increase in processing speed.

## 4. Discussion

Accurate data on species composition and size distribution are essential for uncovering the mechanisms through which environmental change impacts ecological communities. However, reliably and efficiently quantifying these properties at scale remains a major bottleneck in ecological research. In this study, we addressed this challenge by developing a deep learning model that segments, classifies, and counts *Daphnia* species from high-resolution images. To our knowledge, this is the first implementation of a YOLO model specifically designed for automated identification and quantification of zooplankton species in their community context. The model achieved strong performance, with an mAP@50 of 0.899 and mAP@50–95 of 0.659 and a 230-fold speedup with moderate non-GPU hardware compared to manual annotation, reflecting both accurate segmentation and classification across the different classes. Importantly, pixel-precise segmentation enables accurate measurement of individual body sizes, supporting robust analysis of shifts in size distributions, something that previous methods, such as manual stereomicroscope-based measurements or lower-resolution image analysis tools, often struggled to achieve due to limitations in precision or throughput.

Compared to other deep learning models in ecological research and monitoring (see Table 2), our approach integrates object detection, instance segmentation, and species-level classification for freshwater zooplankton, enabling detailed and scalable community-level analysis. While recent methods employing key-point detection (Zhang et al., 2024) or custom CNNs (Karatzas et al., 2020; Rutter et al., 2024) achieve high accuracy for automated size measurement of individual organisms, they are either tailored to single-species images or lack the capacity for simultaneous segmentation and classification within diverse, mixed communities. By contrast, our framework combines these capabilities, supporting comprehensive and high-throughput assessment of both community composition and size structure in complex, multi-species samples.

**Table 2:** Comparison of deep learning models used in ecological and aquatic research, highlighting their capabilities in object detection, instance segmentation, species identification and size measurement, along with their intended purpose.

| Models | Object detection | Instance segmentation | Species Identification | Size Measurement | Purpose | Studies |
| --- | --- | --- | --- | --- | --- | --- |
| Custom CNNs | Yes | No | No | Yes (malformation /size change detection) | Predict malformations in *Daphnia magna* due to ENM exposure using deep learning | (Karatzas et al., 2020) |
| YOLO-EP | Yes | No | No | No | Detect small egg targets in UAV imagery for early pest control in rice agriculture | (Huang et al., 2023) |
| Custom ResNet | Yes (Key-point detection) | No | Yes | Yes | Automated measurement of zooplankton body size in marine taxa using deep learning key point detection | (Zhang et al., 2024) |
| LeDaphNet | Partial (Image Classification) | No | No | Partial (discrete size classes, not continuous) | Semi-automated counting of *D. magna* using a CNN to improve efficiency over manual methods | (Rutter et al., 2024) |
| crawler-wally | Yes | No | Yes | No | Real-time species classification and counting from deep-sea video for autonomous ecological monitoring | (Ortenzi et al, 2024) |
| HydroSpot-YOLO | Yes | No | No | No | Monitor spatial distribution of invasive aquatic plants (*Eichhornia crassipes*) as ecological health indicators | (Ji et al., 2025) |
| **DaphnAI** | **Yes** | **Yes** | **Yes** | **Yes** | **Automate identification, segmentation, for freshwater zooplankton to study community structure and size distribution** | **This study** |

The framework developed offers a powerful tool for ecological research by enabling high-resolution assessment of zooplankton community composition and population size structure with significantly reduced labor and time requirements. Conventional zooplankton community studies typically require the manual inspection of thousands of individuals under a stereomicroscope to detect experimental treatment effects or track population dynamics over time in monitoring projects (Ha and Hanazato, 2009; Pantel et al., 2015; Van Doorslaer et al., 2010). In addition, demographic analyses and biomass estimates of specific populations require the measurement of at least 30, but preferentially >100 individuals per species per sample. In contrast, our approach allows for the identification and measurement of substantially larger numbers of individuals of a fraction of the cost and effort, greatly improving the depth, efficiency, and robustness of ecological data collection. For example, by rapidly quantifying proxies for individual body size (e.g., surface area, width, and length) for large numbers of individuals, our method strongly enhances the ability to detect shifts in size distributions within populations and communities, providing deeper insights the ecological causes and consequences. Changes in zooplankton size structure can indicate shifts in predation regimes, alter grazing pressure on phytoplankton, affect the energy transfer through the aquatic food web, and influence nutrient cycling phytoplankton (Cyr and Downing, 1988; Lampert, 1987; Peters and Downing, 1984; To et al., 2024). Such size-based changes also impact overall ecosystem respiration, linking population or community dynamics to broader biogeochemical processes (Yvon-Durocher et al., 2012). While our model does not explicitly detect fine-scale morphological structures such as *Daphnia* neckteeth or helmets, it enables the extraction of circularity and perimeter as proxies for body shape (Beck et al., 2023). Combined with automated object classification, these outputs could potentially lay the groundwork for high-throughput tracking of morphological variation among individuals within and among populations of different experimental treatments. Alternatively, the outputs generated by our model could serve as inputs for training or use of downstream machine learning models designed to enable finer-scale phenotyping, potentially allowing for the automated detection of subtle morphological traits and more detailed analysis of phenotypic diversity (Becker et al., 2022; Karatzas et al., 2020).

Another key advantage of our approach is the high level of consistency and reproducibility that is ensured by employing a standardized, image-based analysis pipeline. Long-term monitoring programs often suffer from inconsistencies introduced by changes in personnel, methodology, or sampling effort, which can obscure ecological trends (Lindenmayer and Likens, 2010; Lovett et al., 2007). Our approach minimizes observer bias and ensures comparability across time, locations, and users. This consistency enables reliable re-analysis of historical datasets alongside new data, supporting experimental designs and monitoring programs that were previously impractical due to time or resource constraints. For example, long term monitoring programs may benefit from our approach by increasing reproducibility and consistency across long-term time series collected by different individuals or sampling campaigns.

Another key advantage of our approach is its affordability and ease of use for deployment and inference. By relying on low-cost flatbed scanners, open-source software, and models that can run on standard local computers, the system is accessible to a wide range of research groups. Additionally, the automated nature of the classification process eliminates the need for personnel training. This practicality supports broader adoption, and also encourages data and model sharing to the community. While model training and hyperparameter tuning requires access to GPU resources, computational demands are largely limited to the model refinement phase. The day-to-day application of the pipeline does not depend on specialized hardware, making it feasible for labs without high-performance infrastructure to perform large-scale image analysis using pretrained models such as ours.

Despite the practical advantages offered by this approach, several limitations must be acknowledged. Like many neural networks, YOLO models function as a “black box,” producing outputs while obscuring the internal decision-making process (Sejnowski, 2020). This lack of transparency is a well-recognized issue in the application of deep learning models (Baraniuk et al., 2020; Borowiec et al., 2022). In ecological applications, the opacity of such models poses challenges when biological insights are inferred from model outputs, especially when the features that drive classifications remain unknown (Borowiec et al., 2022). Nevertheless, the concept of the “effective black box” remains valid in practice. While the internal mechanics of a model may not be interpretable, the outputs are sufficiently consistent, accurate, and reproducible to be useful for empirical research. In high-throughput contexts, such reliability can outweigh the need for full interpretability, particularly when the goal is to automate repetitive measurements across large datasets (Baraniuk et al., 2020).

Another critical limitation is the model’s sensitivity to annotation quality, particularly in classes where boundaries are either subjective or visually ambiguous (Matabos et al., 2017). In our case, the ballooned class was prone to misclassification due to the similarities to *Daphnia* debris or fragmented individuals, and it is therefore not surprising that the model occasionally predicted *Daphnia* debris as ballooned animals (see SFigure 3). Also, the boundary for annotating a *Daphnia* individual as “ballooned” versus another class is inherently subjective. The transition from intact to ballooned morphology is gradual and difficult to define during annotation. A similar limitation was observed in the egg class, which yielded comparatively lower mAP@50–95 scores. This is primarily attributable to frequent occlusion of eggs within adult *Daphnia* bodies, which limits their visibility (see SFigure 4). As a result, both the annotator and the model struggled to consistently identify and delineate these objects as distinct instances.

In terms of classification performance, morphologically distinct taxa such as *S. vetulus*, *D. galeata*, adult *D. pulex* and copepods were accurately classified and segmented, yet *D. pulex* juveniles were misclassified as *D. galeata (see* Figure 6). This pattern indicates that object size may be a key driver in the classification of *D. pulex*, with the model implicitly associating bigger individuals with that class. While size is a relevant phenotypic trait, relying too heavily on it hinders accurate class separation, especially in experiments which include *Daphnia* species across different life stages. Future attempts might explore classifying all juveniles under a single category while differentiating species only among adults or implementing image augmentation techniques such as random scaling during training could help reduce the model’s reliance on absolute size. This would encourage the network to focus on morphological features other than body size for species classification.

## Conclusion

This study presents a proof of concept as an automated assessment of zooplankton community composition and population size structure tool. The ability to efficiently quantify, classify species and measure their size distributions overcomes critical bottlenecks in ecological monitoring and research. It allows researchers to collect large volumes of high-resolution data that would otherwise require intensive manual effort. This capability makes it possible to design more complex experiments and carry out long-term ecological monitoring with greater efficiency. Future work would focus on bringing computer vision techniques closer to researchers, regardless of coding expertise, by prioritizing user-friendliness across all stages of the workflow. This includes simplifying tasks such as project setup, data annotation, model training, evaluation, and deployment. Such tools set the foundation for a new generation of ecological studies. They provide the precision needed to capture individual-level variation while maintaining the scalability required for population-level assessments.

## Declarations

## Acknowledgments

For training the model the HPC-cluster Hummel-2 at University of Hamburg was used. The cluster was funded by Deutsche Forschungsgemeinschaft (DFG, German Research Foundation) – 498394658. We would like to thank Lyn Westphal for linguistic help.

## Authors Contributions

**SM**, **SAJD**, and **KAO** conceived and designed the study. **ST**, **SAJD**, and **VvS** developed and optimized the scanning procedure; **SM** carried out all data preprocessing and curation, model development and formal analysis. **VvS** provided the raw image data used for this dataset. **SM** wrote the first draft of the manuscript. All authors contributed to reviewing and editing the manuscript and approved the final version. **SAJD** and **KAO** supervised the project.

## Ethical Approval

*This is not applicable*.

## Consent to Participate

*This is not applicable*.

## Consent to Publish

*This is not applicable*.

## Data Availability

All data and code supporting the findings of this study are publicly available at Zenodo under the following DOI: 10.5281/zenodo.15719995.

Equations

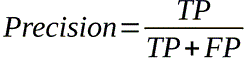

*Equation 1: Precision, calculated as the number of true positives (TP) divided by the total number of predicted positives including false positives (FP)*.

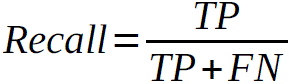

*Equation 2: Recall, calculated as the number of true positives divided by the total number of actual positives including false negatives (FN)*.

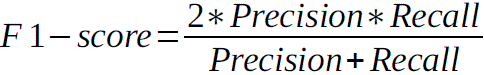

*Equation 3: F1-score, the harmonic mean of Precision and Recall, balancing false positives and false negatives*.

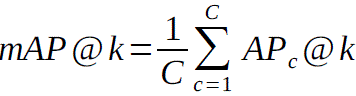

*Equation 4: Mean Average Precision at IoU threshold k (mAP@k) measures the average detection accuracy across all classes, where C is the number of classes and AP_c_@k is the Average Precision for class c at IoU threshold k*.

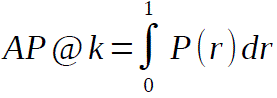

*Equation 5: Average Precision (AP), calculated as the area under the precision–recall curve at IoU threshold k*.

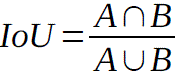

*Equation 6: Intersection over Union (IoU) measures the overlap between the predicted segmentation (A) and the ground truth (B). It is defined as the ratio of their intersection over their union*.

## Supplement

**SFigure 1:**
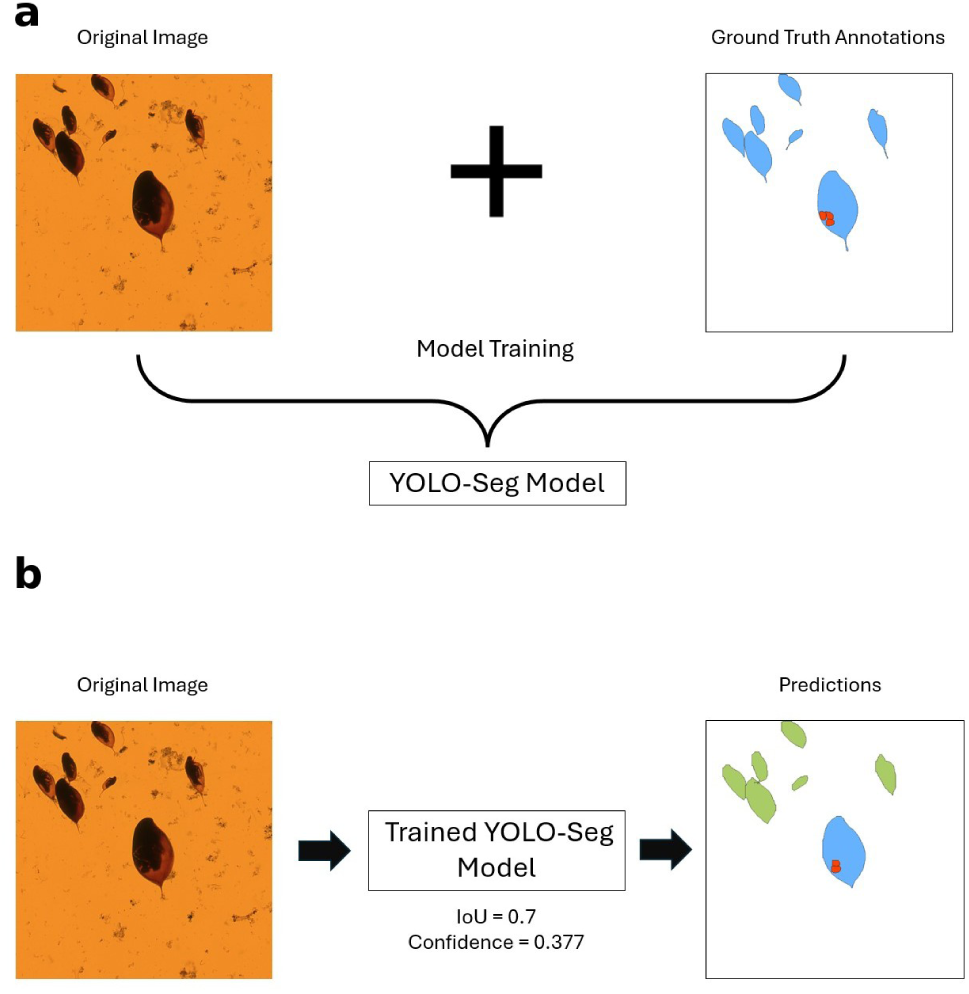
YOLO-Seg training and inference pipeline. a) The original image and its corresponding ground truth annotations, where different colors represent different biological classes.b) Once trained, the model predicts instance masks from input images.

**SFigure 2:**
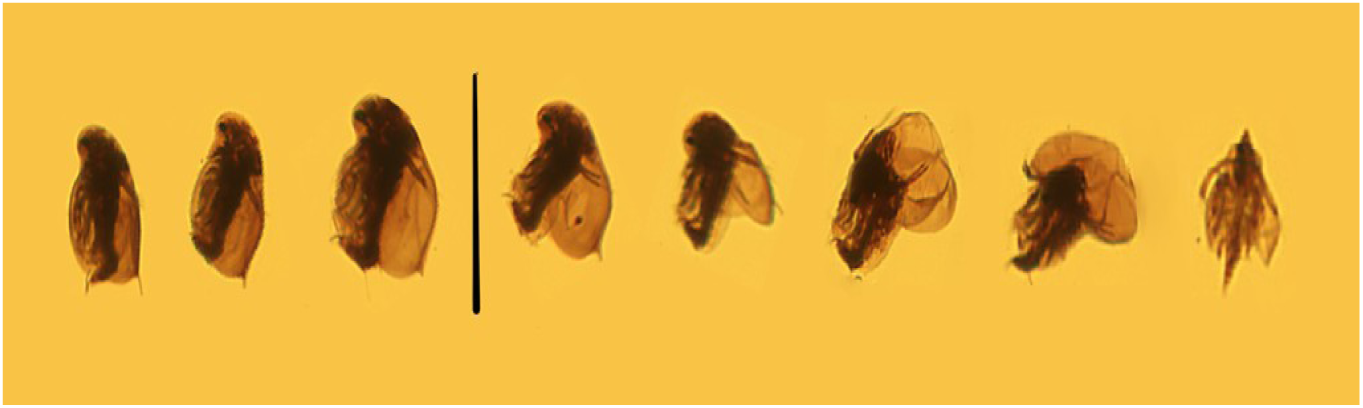
Gradual morphological distortion of Daphnia due to preservation-induced ballooning. The image shows representative individuals arranged from left (normal morphology) to right (severely distorted). The black vertical line indicates the threshold beyond which individuals were classified as “ballooned,” based on the loss of key anatomical features such as the head, tail spine, and overall body curvature.

**SFigure 3:**
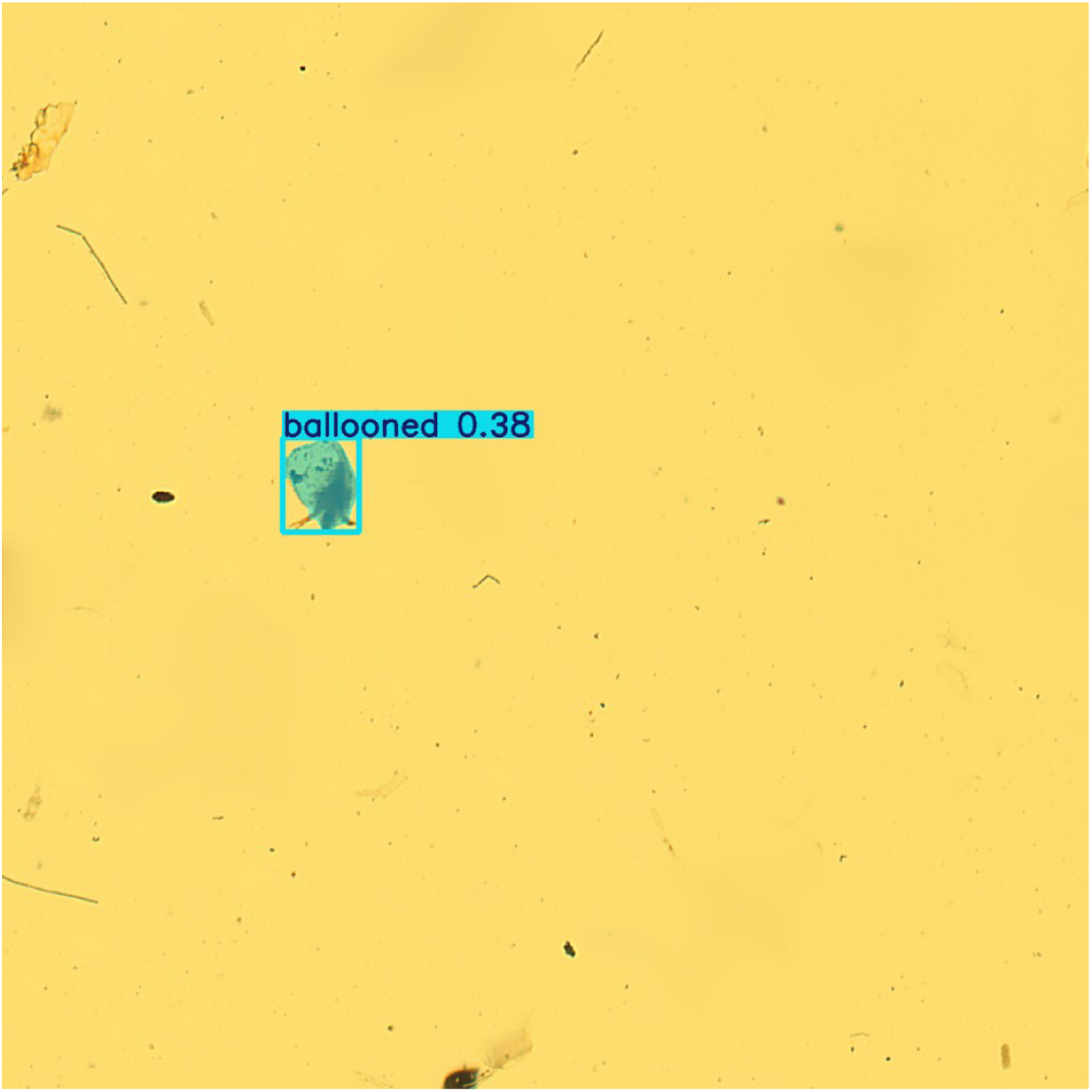
Example of a Daphnia fragment being misclassified as a ballooned individual.

**SFigure 4:**
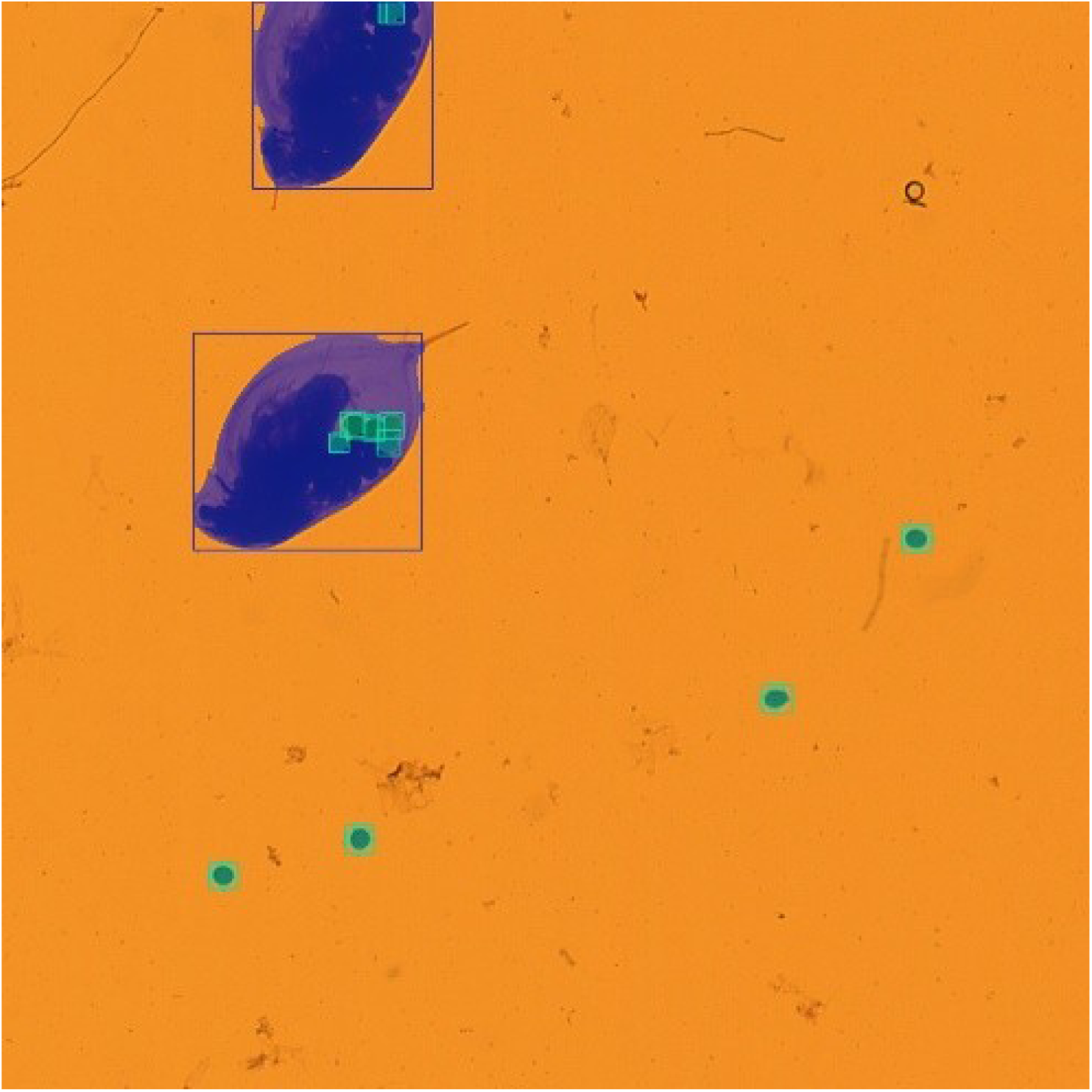
Detection and segmentation of eggs (green masks) within Daphnia individuals (blue boxes). Eggs that are fully enclosed within the body of the Daphnia, makes their boundaries ambiguous. This poses challenges for both model predictions and human annotations, leading to inconsistent detection and reduced segmentation performance.

